# Dissociable neuronal modules in the orbitofrontal cortex representing single and multi-attribute choice tasks

**DOI:** 10.1101/2022.07.02.498538

**Authors:** Chengze Xu, Kuntan Ni, Xinying Cai

## Abstract

An important component of biological intelligence lies in the capacity to learn and execute various cognitive tasks. This ability may be facilitated by a neural system comprising functionally dissociable neuronal modules. Within the orbitofrontal cortex (OFC) lies a neural circuit that supports economic decision-making across diverse contexts. To investigate the functional specialization of this circuit, we compared the neural activity of OFC neurons in rhesus monkeys during multi-attribute choice (MC) and single-attribute choice (SC) tasks. In MC, the monkeys made subjective tradeoffs between competing attributes, whereas in SC, choices were deterministic based on a single attribute’s value. Neuronally, in MC, OFC neurons primarily encoded offer value, chosen value, and choice in goods space. Upon transitioning from MC to SC, a significant portion of MC-modulated neurons disengaged, while a separate set of neurons emerged to encode SC-related variables in a spatial reference frame. Notably, we observed the clustering of value-encoding neurons in MC but not SC. In essence, our findings suggest that choice tasks involving different mental processes are represented by dissociable neuronal modules within the OFC.

## Introduction

The primate brain can handle various cognitive tasks and flexibly switch among them, yet the precise neural mechanisms remain elusive. One plausible explanation is the presence of functionally dissociable neuronal modules, with each module predominantly activated to meet specific task demands (Yang et al., 2019a; Yang et al., 2019b). An advantage of having functionally dissociable modules is that structural knowledge of a common meta-task can be abstracted, enabling its generalization to different instances (Behrens et al., 2018). In this regard, although seemingly paradoxical, circuit-level functional specialization may facilitate task generalization.

Numerous experimental and theoretical studies have illustrated the flexible coding capabilities of the OFC across various choice contexts (Cai and Padoa-Schioppa, 2019; Conen and Padoa-Schioppa, 2019; Kobayashi et al., 2010; Padoa-Schioppa, 2009; Padoa-Schioppa and Assad, 2008; Rustichini et al., 2017; Tremblay and Schultz, 1999; Xie and Padoa-Schioppa, 2016). However, the putative choice circuitry within the OFC has seldom been investigated under paradigms involving different choice tasks, leaving it unclear whether the OFC houses functionally dissociable modules for such diverse demands.

Moreover, while functional specialization has been proposed as one of the fundamental organizational principles of the brain (Kanwisher, 2010), there is limited understanding of task representations at the neuronal level, as existing studies have primarily focused on analyzing specialization across brain regions (Connolly et al., 2003; Conway et al., 2007; Downing et al., 2001; Gavrilov et al., 2017; Kanwisher et al., 1997; Tsao et al., 2006; Zeki, 1978). Neuronal-level functional specialization is particularly relevant to the prefrontal cortex (PFC) due to its cognitive versatility. We hypothesize that task specialization in the PFC may occur at a finer resolution than suggested by previous studies (O’Reilly, 2010; Riley et al., 2017). Therefore, cross-task single-neuron analysis is essential for studying the functional specialization of the PFC concerning higher cognitive functions.

To investigate the functional specialization of OFC neuronal ensembles in value-based decisions, we conducted a study in which monkeys switched between two different choice tasks: a multi-attribute choice (MC) task and a single-attribute choice (SC) task. In the MC task, the animal chose between two juices of varying amounts, and its choice pattern exhibited a subjective tradeoff between juice taste and quantity. In the SC task, the animal chose between the same juice in different quantities, with the choice directed towards the larger quantity offer deterministically. These distinct choice patterns suggested different mental operations under the two tasks. Are these tasks supported by dissociable neuronal modules, or do they share the same population of neurons?

To investigate this, we recorded the activity of the same neurons across both tasks. In the MC task, OFC neurons predominantly encoded the same parameters as previously reported: offer value, chosen value, and chosen juice. However, upon switching to the SC task, a significant portion of the MC-related neurons disengaged, while a new set of neurons emerged to encode SC-related variables. Interestingly, we observed that neurons encoding the same value-related variable tended to cluster anatomically, but this clustering was only evident in the MC task.

By contrasting the neuronal representations between MC and SC tasks, our study provides physiological evidence supporting the existence of dissociable neuronal modules in the OFC, each representing decisions involving different mental operations.

## Results

### Behavior

From the behavioral perspective, economic choice is characterized as entailing a tradeoff among competing attributes, and preference towards these attributes is reflected in the idiosyncratic tradeoff patterns across decision-makers as well as for the same decision-maker under different circumstances. In our study, we investigated how neuronal representations in the OFC adapt when animals transition from multi-attribute choice (MC) blocks (A:B) to single-attribute choice (SC) blocks (B:B and A:A) (see **Fig. 1a**). Each experimental session comprised four blocks of trials arranged in the sequence of B:B, (A:B)_1_, (A:B)_2_, A:A (refer to **Methods**). Different pairs of juices were presented in various experimental sessions, with the same two juices consistently used within a single session. To facilitate a matching analysis similar to that for the A:B block, one of the two offers in B:B blocks, where the same juice was presented for each offer, was denoted as lowercase “a”. Likewise, for A:A blocks, one offer was designated as lowercase “b” (see **Fig. 1b**). In subsequent analyses, when applicable, (A:B)_1_ and (A:B)_2_ blocks are collectively referred to as the A:B task, while B:B and A:A blocks are collectively referred to as the X:X task.

**Figure 1.**
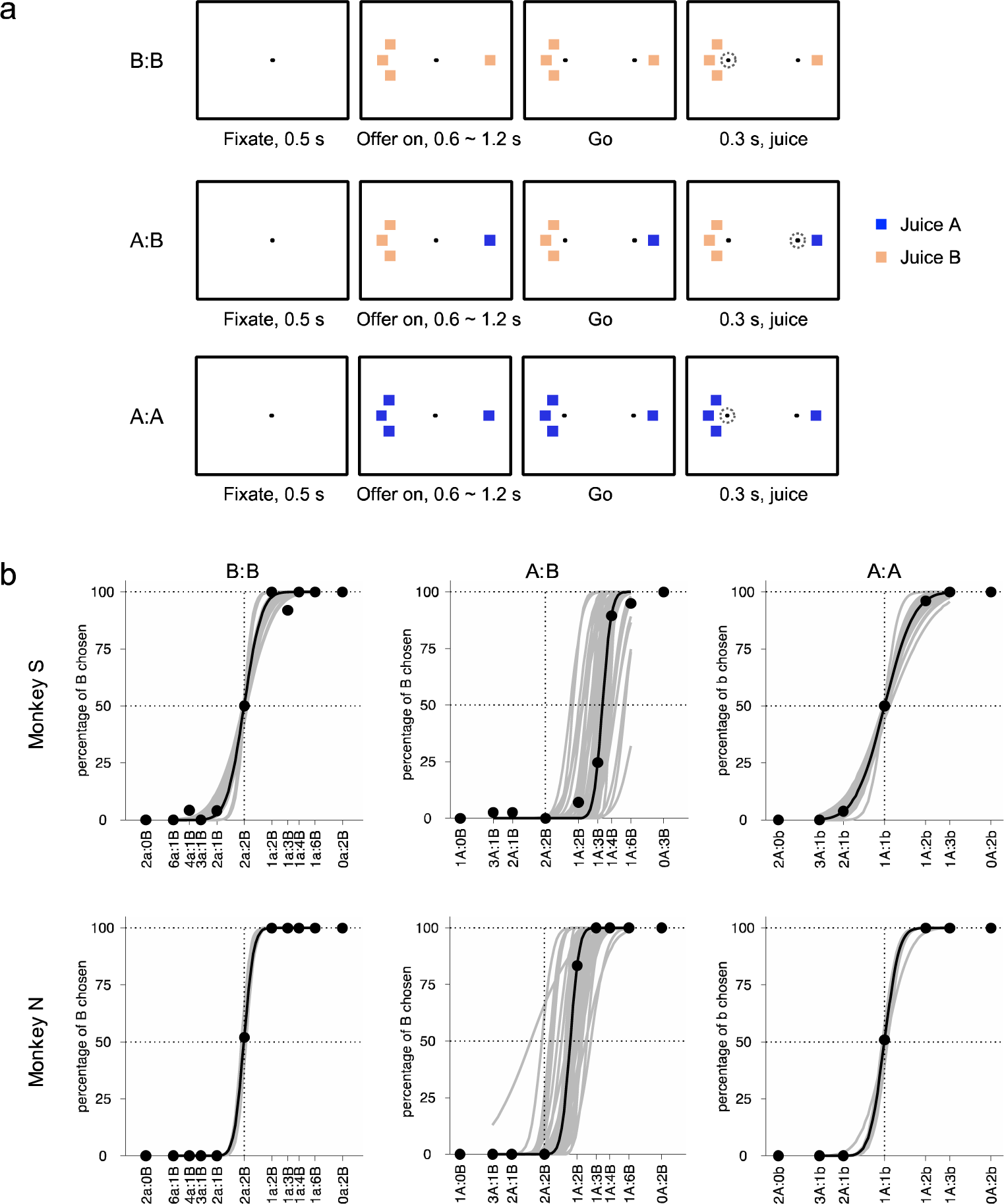
Experimental design and behavioral analysis. (**a**) The same two juices (A and B) were used in one experimental session with juice A the preferred one. One experimental session typically includes 4 blocks of trials in the sequence of B:B, A:B, A:B and A:A. The temporal structure of the task is the same for all blocks. (**b**) Choice patterns, all sessions. For the purpose of performing a matching analysis as that for the A:B blocks, in B:B blocks, one of the two offers was coded as lowercase ‘a’ and in A:A blocks, one of the two offers was coded as lowercase ‘b’. Each gray line indicates the sigmoid fit for one block of trials. The black dots and the black line of sigmoid fit are based on the trial block with the median r value among all blocks.

In the A:B task, both animals consistently exhibited choice relative values significantly larger than one (refer to **Methods, Fig. 1b**, monkey S (44 sessions), ρ = 3.36±0.96 (mean±std), p < 10^-19^, t-test; monkey N (59 sessions), ρ = 1.75±0.35 (mean±std), p < 10^-23^, t-test). Conversely, in the X:X task, there were no significant differences in choice relative values from one for either animal (**Fig. 1b**, monkey S (46 sessions), ρ = 1.005±0.029 (mean±std), p = 0.23, t-test; monkey N (51 sessions), ρ = 0.999±0.023 (mean±std), p = 0.66, t-test). Additionally, in the X:X task, the animals’ choices were predominantly influenced by offer quantity, as evidenced by their high accuracy in selecting the offer with the larger quantity (**Fig. 1b**, monkey S: B:B condition (23 sessions), 97.7%±1.7%, A:A condition (23 sessions), 98.6%±1.2%; monkey N: B:B condition (34 sessions), 99.97%±0.16%, A:A condition (17 sessions), 100%±0%).

The contrast in choice patterns between the A:B and X:X tasks suggests that the animals were engaged in different mental operations. In the A:B task, the animals’ choices were influenced by comparing the subjective values that integrate both juice taste and quantity. Meanwhile, in the X:X task, the choice was solely based on comparing the quantity of the same juice. Subsequently, we investigated how these two different mental operations are represented in the OFC.

### Neuronal encoding in MC and SC

Our encoding analysis was conducted on 2029 cells in the A:B task (938 from monkey S; 1091 from monkey N) and 1964 cells in the X:X task (912 from monkey S; 1052 from monkey N) (**Fig. 2**; see **Methods**). We first examined the encoding of task-related variables in both the A:B and X:X tasks. In the A:B task, OFC neurons predominantly encoded the same set of variables as previously reported. However, most of these neurons disengaged when the animal transitioned to the X:X task.

**Figure 2.**
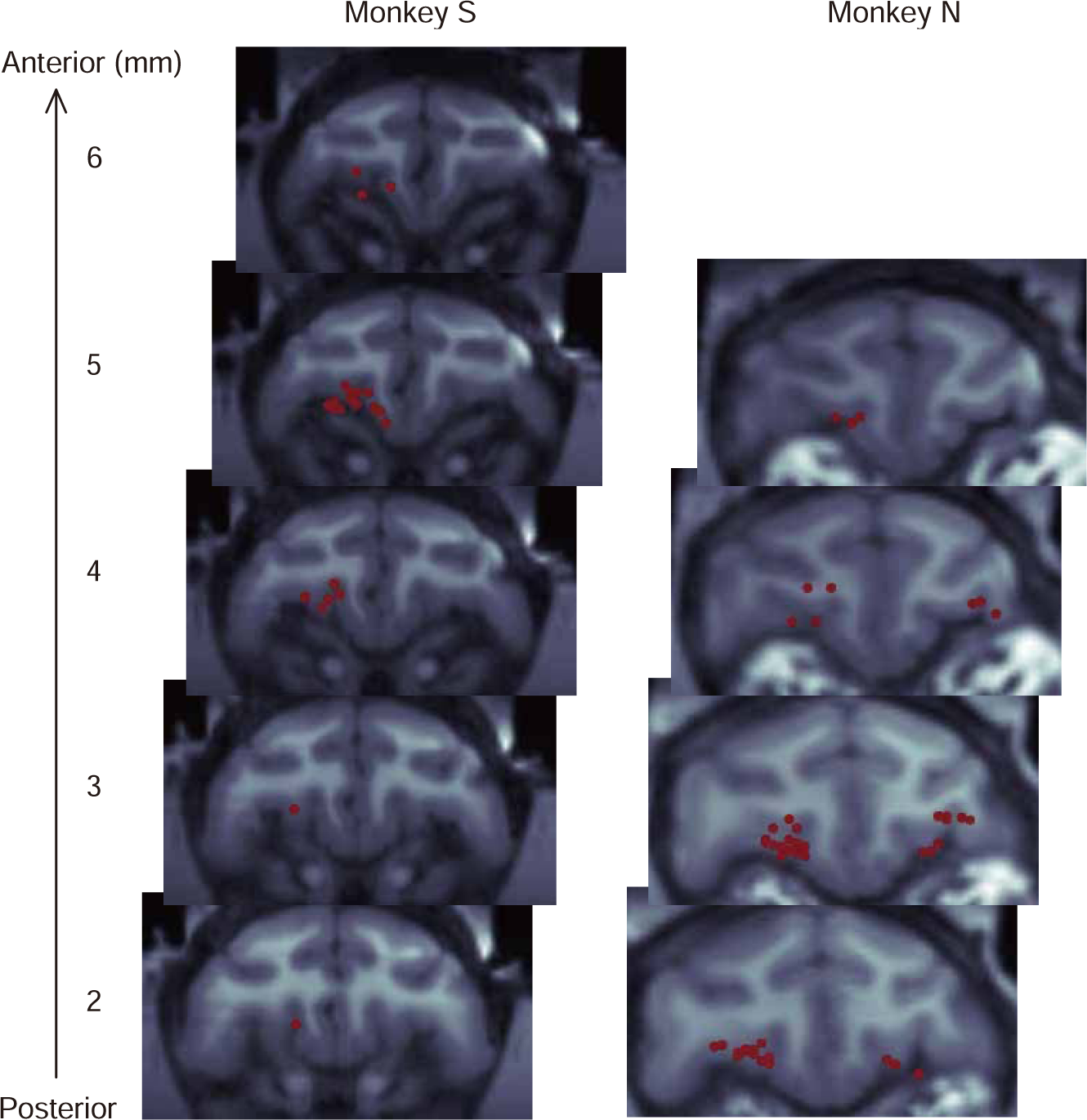
MRI reconstruction of recording sites. For monkey S, the recording covered 5 coronal (1 mm spacing) from 2 to 6 mm anterior to the head of the genu, while for monkey N, the recording covered 4 coronal slices (1 mm spacing) from 2 to 5 mm anterior to the head of the genu. The recording was performed in the central OFC, located between the medial orbital sulcus and the lateral orbital sulcus. Each red dot indicates one recording site on 16-channel V/S probes (Plexon Inc) or the tip of individual tungsten electrodes tungsten electrodes (Frederick Haer).

Here, we present five example neurons recorded during the A:B task (**Fig. 3**). One neuron (**Fig. 3a**) encoded the offer value of juice A, while another neuron (**Fig. 3b**) encoded the offer value of juice B. A third neuron (**Fig. 3c**) encoded the chosen value, while a fourth neuron (**Fig. 3d**) encoded the chosen juice. Additionally, a fifth neuron encoded the number of square symbols on the contralateral hemifield (Symbol #_contra_). Importantly, few of these neurons encoded task-related variables during the X:X task.

**Figure 3.**
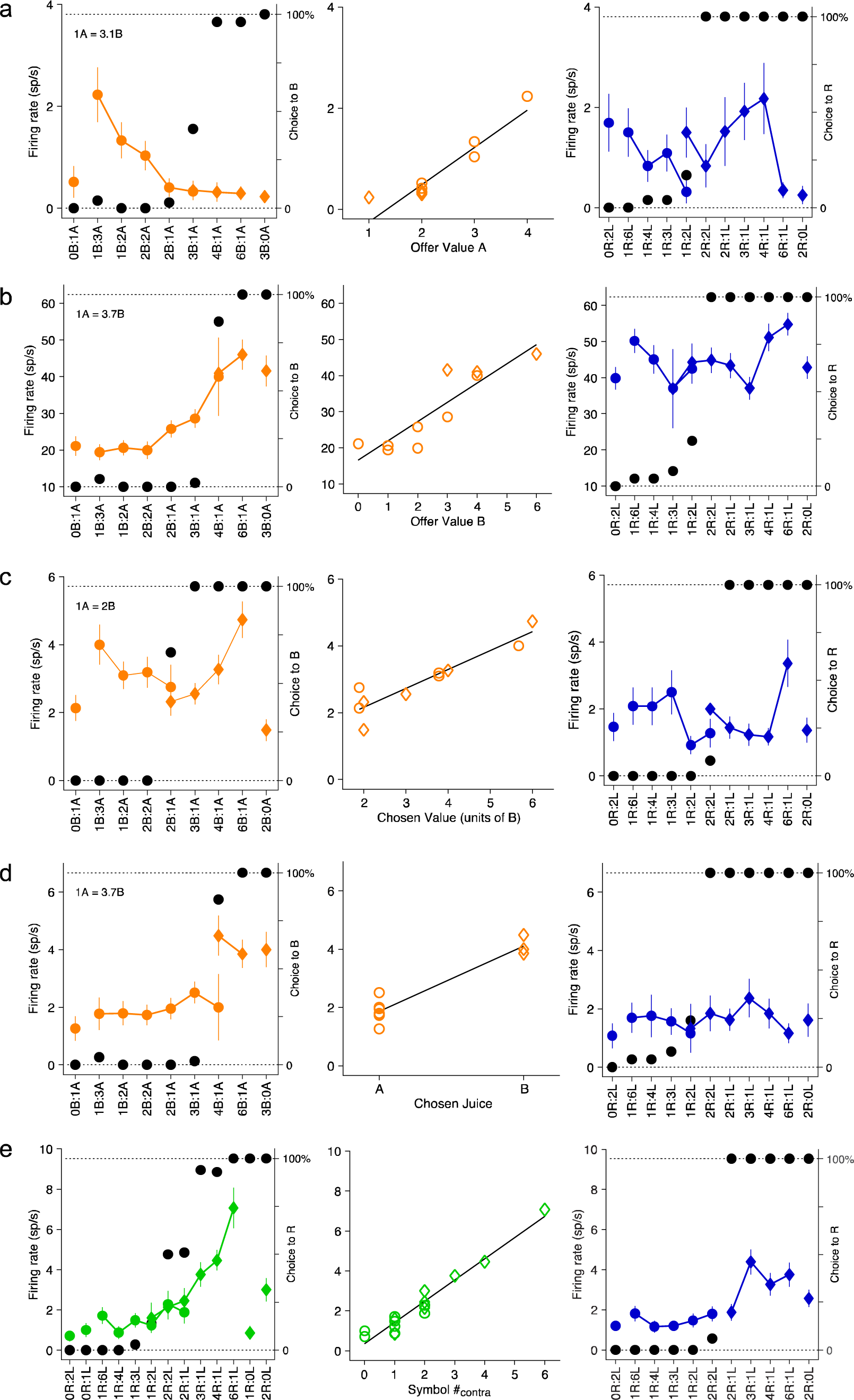
Example neurons encoding task-related variables in MC. In **a-d**, the left panel shows the activity of the cell recorded in the A:B task; The x axis represents different offer types ranked by ratio #B:#A. Black symbols represent the percentage of ‘‘B’’ choices. Orange symbols represent the neuronal firing rate (circles and diamonds for choices of juice A and juice B). In **e**, the left panel shows the activity of the cell recorded in the A:B task; The x axis represents different offer types ranked by ratio #R:#L. Black symbols represent the percentage of ‘‘R’’ choices. Green symbols represent the neuronal firing rate (circles and diamonds stand for choices to the left and right). In all figures, the middle panel shows the activity of the neuron plotted against the encoded variable; The black line is derived from a linear regression. The right panel shows the activity of the same cell but recorded in the adjacent B:B block. The x axis represents different reward quantity combinations ranked by ratio #R:#L. Black symbols represent the percentage of ‘‘R’’ choices. Blue symbols represent the neuronal firing rate (circles and diamonds for choices to the left and right). (**a**) Cell encoding Offer Value A (R^2^ = 0.88, p < 10^-4^). This cell did not encode any defined task-related variables (Table S1) in the B:B task. (**b**) Cell encoding Offer Value B (R^2^ = 0.80, p < 10^-3^). This cell encoded the Chosen Quantity in the B:B task (R^2^ = 0.37, p < 0.05). (**c**) Cell encoding Chosen Value (R^2^ = 0.86, p < 10^-3^). This cell did not encode any defined task-related variables in the B:B task. (**d**) Cell encoding Chosen Juice (R^2^ = 0.91, p < 10^-4^). This cell did not encode any defined task-related variables in the B:B task. (**e**) Cell encoding Symbol #_contra_ (R^2^ = 0.94, p < 10^-9^). This cell encoded Offer Quantity_contra_ in the B:B task (R^2^ = 0.57, p < 0.05). Firing rates shown here are from the post-offer time window. For these neurons, the numbers of trials in the A:B and B:B block were as follows: (**a**) 269, 265, (**b**) 290, 284, (**c**) 308, 273, (**d**) 290, 284, (**e**) 285, 342.

Conversely, many neurons encoding task-related variables in the X:X task became untuned in the A:B task. Here, we showcase three example neurons (**Fig. 4**). One neuron (**Fig. 4a**) encoded the quantity of the contralateral offer (Offer Quantitycontra), while another neuron (**Fig. 4b**) encoded the quantity of the chosen offer (Chosen Quantity). A third neuron (**Fig. 4c**) encoded the location of the chosen offer (Chosen Side). Strikingly, all of these neurons exhibited untuned activity in the A:B task. For population analyses, our approach was as follows. In the A:B task, we defined a “trial-type” by considering two offers, their spatial arrangement, and the choice made (e.g., [1A+:3B-, A]). Here, “+” indicated an offer located on the left hemifield, while “-” indicated an offer on the right hemifield. Conversely, in the X:X task, since only one type of juice was presented, and the animals consistently chose the offer with a larger quantity, the “trial-type” was defined by two offers and the location of the larger quantity offer (e.g., [1B+:3B-]). Our neuronal analysis focused on the post-offer window, spanning 100 to 600 ms after offer presentation. For each neuron, we conducted a one-way ANOVA (p<0.01) across trial types. Neurons meeting this criterion were identified as task-related and included in subsequent variable selection analysis, comprising 350 out of 2029 cells (17.2%) in the A:B task and 157 out of 1964 cells (8%) in the X:X task.

**Figure 4.**
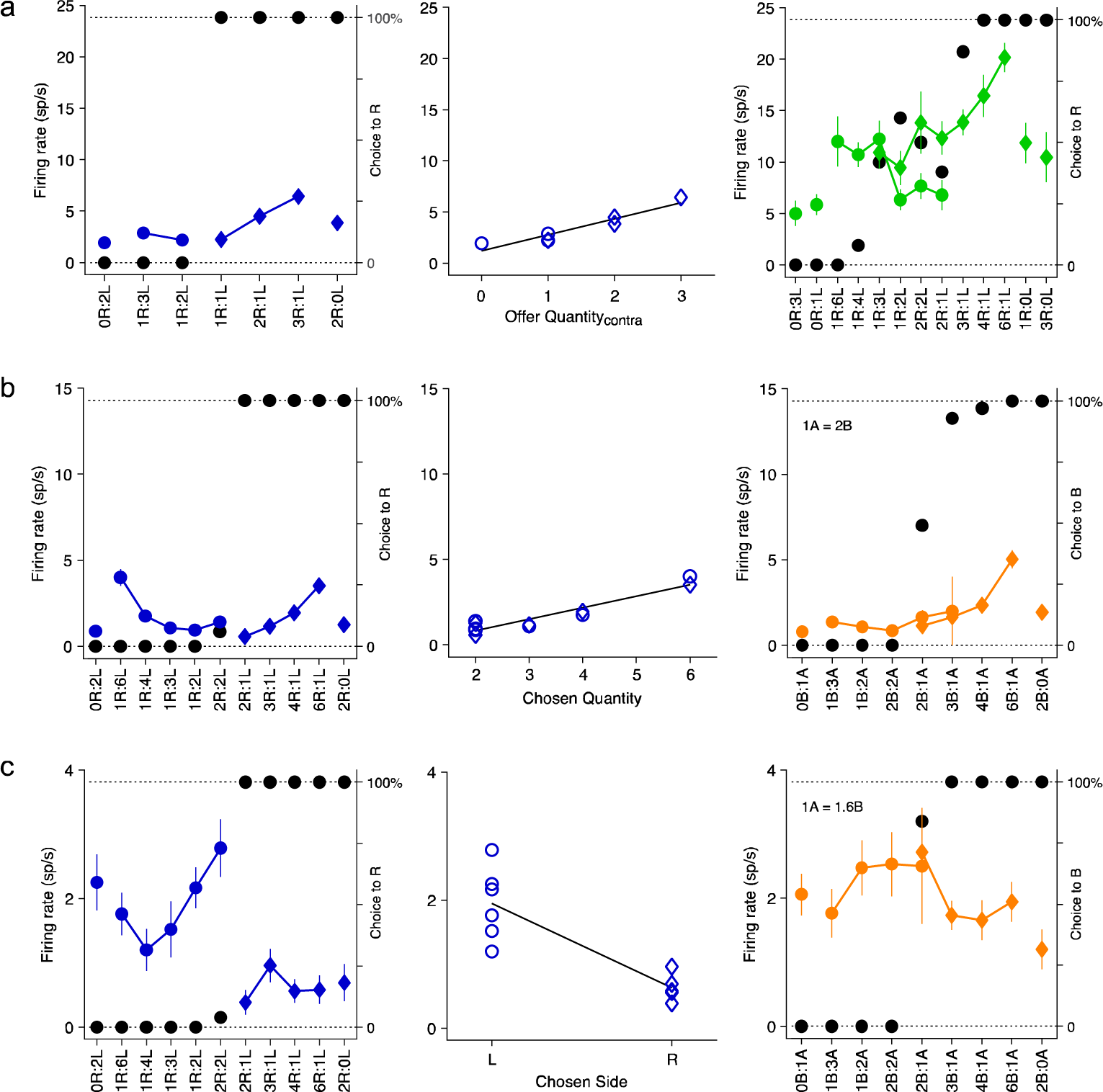
Example neurons encoding task-related variables in the SC. The left panel shows the activity of the cell recorded in the A:A or B:B block; The x axis represents different reward quantity combinations ranked by ratio #R:#L; Black symbols represent the percentage of ‘‘R’’ choices; Blue symbols represent the neuronal firing rate (circles and diamonds for choices to the left and right). The middle panel shows the activity of the neuron plotted against the encoded variable; The black line is derived from a linear regression. The right panel shows the activity of the cell recorded in the adjacent A:B block of the same session. The x axis represents different offer types ranked by ratio #B:#A. Black symbols represent the percentage of ‘‘B’’ choices. Orange symbols represent the neuronal firing rate (circles and diamonds for choices of juice A and juice B). (**a**) Cell encoding Offer Quantity_contra_ (R^2^ = 0.89, p < 0.01). This cell encoded Symbol #_contra_ in the A:B task (R^2^ = 0.40, p < 0.05). (**b**) Cell encoding Chosen Quantity (R^2^ = 0.89, p < 10^-4^). This cell did not encode any defined task-related variables in the A:B task (Table S1). (**c**) Cell encoding Chosen Side (R^2^ = 0.72, p < 10^-3^). This cell did not encode any defined task-related variables in the A:B task. The firing rates shown here are from the post-offer time window. For these neurons, the numbers of trials in each trial block were as follows: (**a**) 342, 266, (**b**) 369, 249, (**c**) 273, 266.

Each neuronal response that met the analysis of variance criterion was subjected to regression against a set of potential encoding variables (**Table S1**). A variable was deemed to “explain” the response if the regression slope significantly deviated from zero (p < 0.05). Additionally, each linear regression provided an R^2^ value, with R^2^ set to 0 for variables that did not explain the response. The variable with the highest R^2^ value was considered the “best fit” for the neuronal response, and the cell was categorized as untuned if it did not encode any variable (see **Methods**). In the A:B task, 308 out of 350 (88%) task-related responses were explained by at least one of the 11 variables examined in the analysis (**Table S1**). To quantitatively assess the variables that best explained the population, we employed the stepwise selection method. As depicted in **Fig. 5a**, the first four iterations of this procedure selected the variables Offer Value_juice_, Chosen Value, Symbol #_spatial_, and Chosen Juice. Together, these four variables accounted for 92% (284/308) of the responses explained by the 11 variables. Subsequent iterations yielded variables with a marginal explanatory power of less than 5%. The best subset method confirmed that these four variables explained more responses than any other subset of four variables. Neurons encoding each of these selected variables (Symbol #_spatial_, Offer Value_juice_, Chosen Value, and Chosen Juice) constituted 14%, 28%, 17.4%, and 21.7% of all task-related neurons (350), respectively (**Fig. 5b**). In the X:X task, 128 out of 157 (82%) task-related responses were explained by at least one of the five variables (**Table S1**). The first three iterations of the stepwise selection procedure selected the variables Chosen Quantity, Offer Quantity_spatial_, and Chosen Side (**Fig. 5c**). Together, these three variables explained 98% (125/128) of the responses explained by the five variables (**Fig. 5c**). The best subset method also selected these three variables. Neurons encoding each of the three selected variables (Offer Quantity_spatial_, Chosen Quantity, and Chosen Side) constituted 29.3%, 41.4%, and 8.9% of all task-related neurons (157), respectively (**Fig. 5d**).

**Figure 5.**
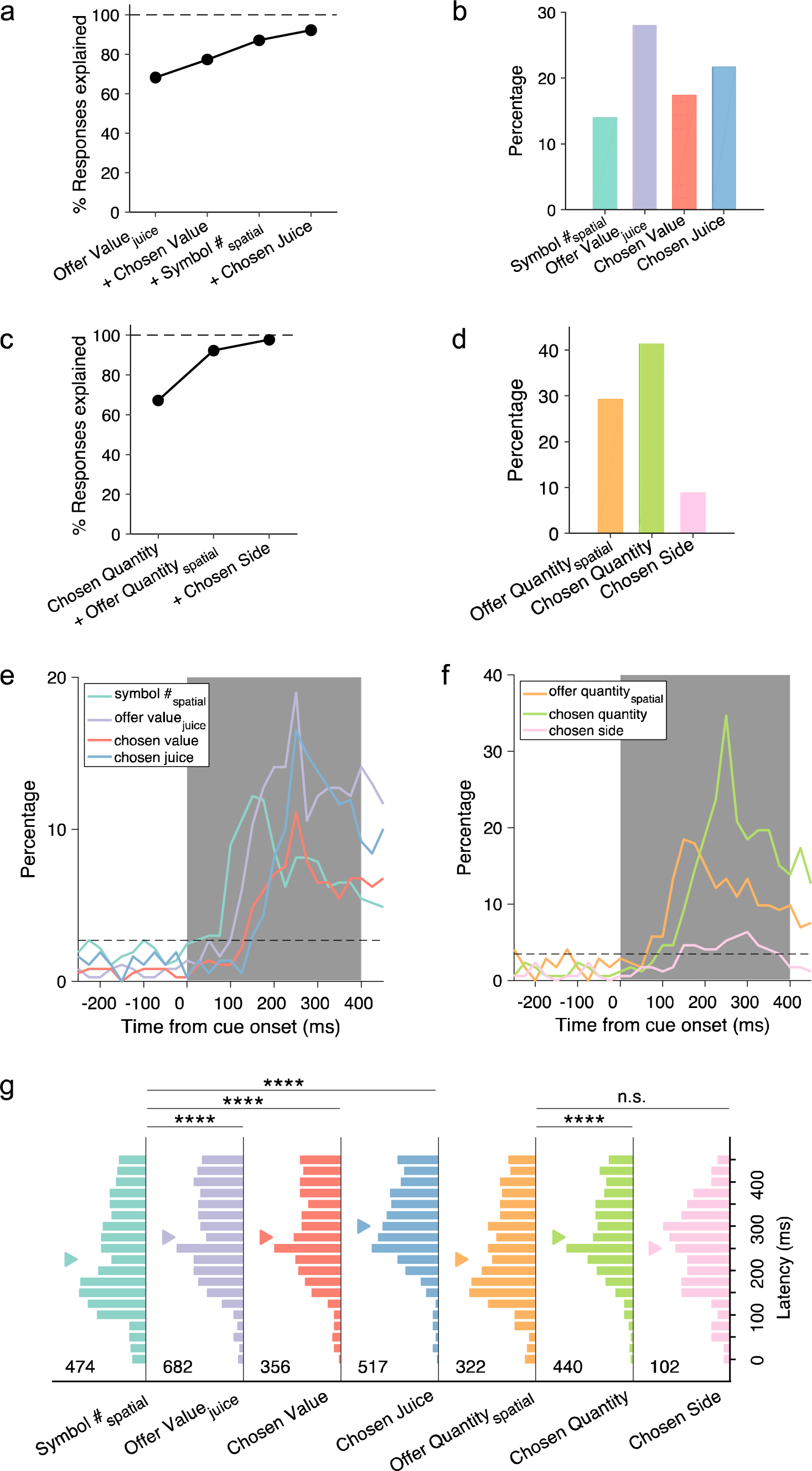
Encoding of task-related variables. (**a**) Stepwise selection, percentage of explained responses. The figure illustrates the percentage of responses explained at subsequent iterations of the stepwise procedure (“+” on the abscissa indicates inclusion of the variable. In this plot, the “100” dotted line represents the total number of responses explained overall by the 11 variables examined for the A:B task (Table S1). The four variables Offer Value_juice_, Chosen Value, Symbol #_spatial_ and Chosen Juice collectively explained 92% of the responses explained by the 11 variables. (**b**) Percentage of neurons encoding each selected variable in the A:B task. (**c**) Same as in (**a**) but for X:X task. The three variables Chosen Quantity, Offer Quantity_spatial_ and Chosen Side explained 98% of the responses explained by the 5 variables examined for the X:X task (Table S1). (**d**) Percentage of neurons encoding each selected variable in the X:X task. (**e**) Time course of percentage of encoded variables in the A:B task. Dotted horizontal line indicates the percentage significantly above the chance level of 1% (binomial test, p < 0.05). Solid vertical line indicates time of offer on. (**f**) Same as in (**e**) but for the encoded variables in X:X task. (**g**) Distribution of encoding latency for encoded variables in A:B and X:X task. Number in each subplot indicates the number of responses encoding the corresponding variable. Difference in latency distribution between Symbol #_spatial_ and other variables in the A:B task was tested using the Kolmogorov–Smirnov test (****p < 0.0001); Difference in encoding latency distribution between Offer Quantity_spatial_ and other variables in the X:X task was tested using the same test (****p < 0.0001; n.s., not significant).

The identification of the variable Symbol #_spatial_ was previously unreported in similar studies. We hypothesize that it may signify a spatial perceptual signal, given its dominance over the potentially confounding variable Offer value_spatial_. To investigate this hypothesis further, we examined the temporal dynamics of encoded variables. Employing 100 ms time bins with 25 ms steps, we analyzed the activity of each cell over time. **Figure 5e** illustrates the percentage of cells encoding each of the four variables at different time points relative to the offer presentation. Interestingly, in the A:B task, the Symbol #_spatial_ signal appears earlier compared to the other variables. Similarly, in the X:X task, the Offer Quantity_spatial_ signal also demonstrates an earlier onset (**Fig. 5f**).

We then conducted a quantitative analysis of encoding latency (EL). For each variable, we defined the time window during which a neuronal response encoded the variable with the best fit (**Fig. 5e and 5f**) as one instance of EL. We aggregated all these time windows to create a distribution of EL for each variable (**Fig. 5g**). Subsequently, we compared the latency distribution of Symbol #_spatial_ with that of other variables in the A:B task and the latency distribution of Offer Quantity_spatial_ with that of other variables in the X:X task. In the A:B task, the EL for Symbol #_spatial_ (225±5.3 ms, median±ste) was significantly shorter than that for Offer Value_juice_ (275±4.1 ms), Chosen Value (275±5.5 ms), and Chosen Juice (300±4.0 ms) (p < 0.0001, KS tests). In the X:X task, the EL for Offer Quantity (225±5.1 ms, mean±ste) was significantly shorter than that for Chosen Quantity (275.0±4.6 ms, p < 0.0001, KS test), but not significantly different from that for Chosen Side (250±10.1 ms, p = 0.68, KS test). The encoding of symbol numbers in the A:B task, occurring earlier than the encoding of value signals, aligns with the findings of attentional signals being processed in the OFC (McGinty et al., 2016; Xie et al., 2018).

### Dissociable neuronal modules for MC and SC

The results from contrasting single neuron encoding across the A:B and X:X tasks provided initial evidence that these tasks may be represented by separate neuronal modules in the OFC. To further investigate systematically, we examined how the encoding of task-related variables remaps across the two tasks. We established the neuronal remapping across the two adjacent A:B blocks as the control condition, referred to as the (A:B)_1_ → (A:B)_2_ condition. In contrast, the remapping in A:B → B:B and A:B → A:A constituted the experimental condition, collectively referred to as the A:B → X:X condition.

In total, 1078 cells were consistently recorded under the (A:B)_1_ → (A:B)_2_ condition, while 1842 cells were consistently recorded under the A:B → X:X condition. Given that cross-block recording stability significantly influences the assessment of neuronal remapping, we first confirmed that there was no significant difference in the stability of cross-block recordings between the (A:B)_1_ → (A:B)_2_ and A:B → X:X conditions. To do so, we computed Pearson’s correlation for both the distributions of inter-spike intervals (**Fig. S1a**) and the spike waveforms (**Fig. S1b**) of the same neuron, as well as for randomly paired cells across the two adjacent blocks. Subsequently, we calculated the recording stability indices (RSI_isi_ and RSI_waveform_) for each neuron. Following this, we evaluated whether there was a significant difference in RSI between the (A:B)_1_ → (A:B)_2_ and A:B → X:X conditions. The analysis revealed no significant difference for both the RSI_isi_ (**Fig. S1c_i** and **S1c_ii**, p = 0.49, Wilcoxon Rank-Sum Test) and RSI_waveform_ (**Fig. S1c_iii** and **S1c_iv**, p = 0.66, Wilcoxon Rank-Sum Test).

To explore the remapping of cross-task encoding, we first constructed a contingency table based on the results of single-cell encoding in the control condition (**Fig. 6a**). The dataset comprised 281 cells that were modulated in at least one of the two blocks. Our observation revealed that neurons tended to concentrate either on the main diagonal or the rightmost column and bottom row. This suggests that neurons encoding task-related variables in (A:B)_1_ tended to encode the same variable or became untuned in the second block ((A:B)_2_) of the same task. For statistical significance testing, we first computed the odds ratio for each element of the contingency table (**Fig. 6b**). Subsequently, we conducted Fisher’s exact test to evaluate, for each position in the contingency table, whether the cell count deviated from the chance level, assuming that the classifications in the two trial blocks were independent. The p-values in **Fig. 6b** indicate whether cell counts were significantly above (red asterisks, p < 0.05) or below (blue asterisks, p < 0.05) chance. All positions on the main diagonal, except for the{untuned, untuned}pair, exhibited cell counts above chance, supporting persistent coding across two blocks of the same task.

**Figure 6.**
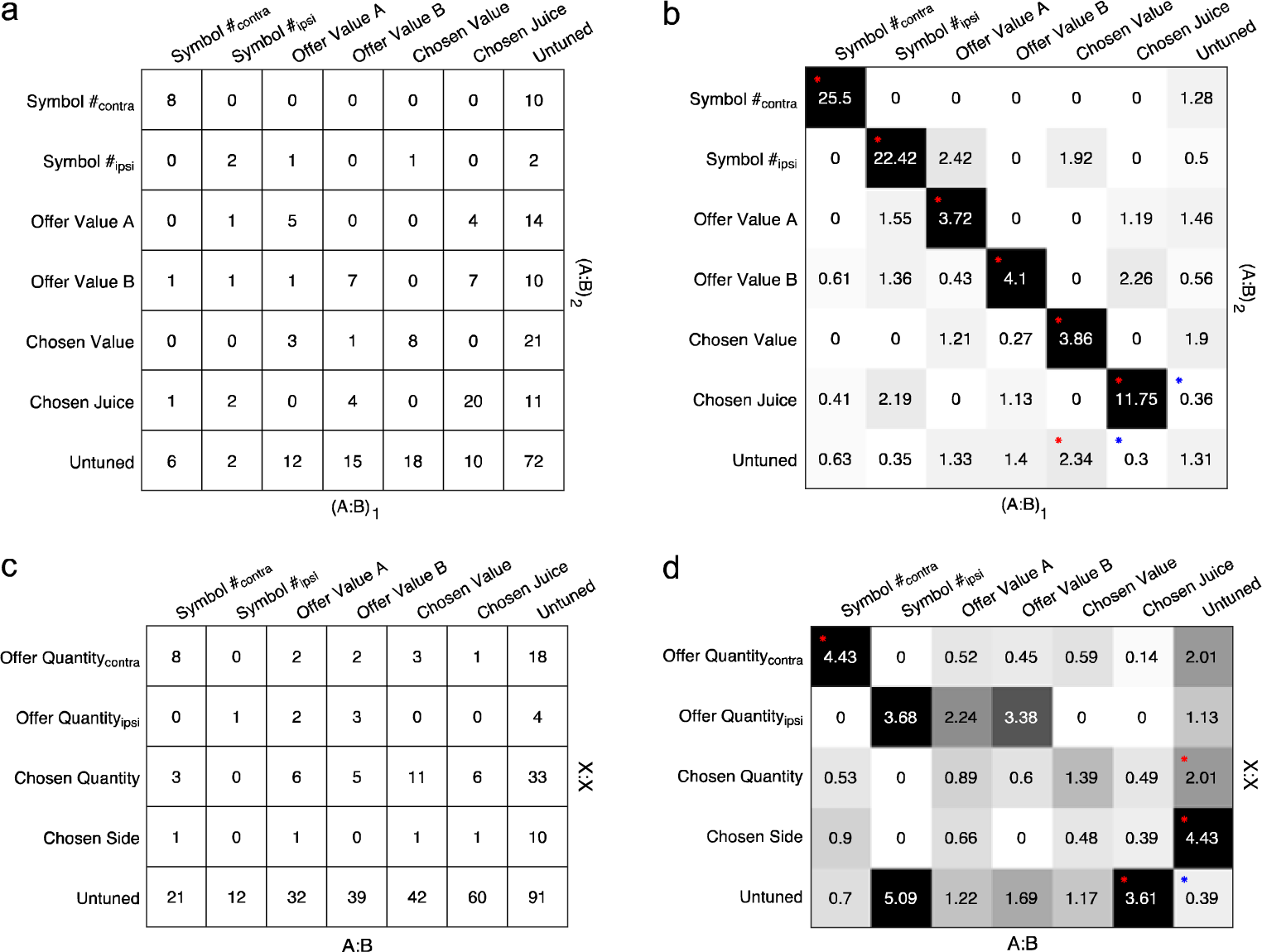
Contrasting classifications across trial blocks. This population analysis was conducted separately for (A:B)_1_→(A:B)_2_ (**a** and **b**) and A:B→X:X (**c** and **d**). (**a**) Contingency table, (A:B)_1_→(A:B)_2_ (*n* = 281). Rows and columns represent, respectively, the classification obtained in the first and the second A:B block. Numbers in the table indicate cell counts. Task-related cells not explained by any variable were classified as untuned. (**b**) Analysis of odds ratios, (A:B)_1_→(A:B)_2_. Numbers in the table and grayscale colors represent the odds ratios obtained for each location in (**a)**. Chance level is 1 and numbers >1 (or <1) indicate that the cell count was above (or below) that expected by chance. For each location, we performed Fisher’s exact test. Red asterisks indicate that the cell count was significantly above chance (p < 0.05) while blue asterisks indicate that the cell count was significantly below chance (p < 0.05). (**c** and **d**) Same as (**a** and **b**) but for A:B→X:X.

Conversely, the contingency table (comprising 419 cells) for the experimental condition displayed a markedly different pattern (**Fig. 6c and 6d**). Apart from the pair {Offer Quantity_contra_, Symbol #_contra_}, there was no consistent mapping in the encoding of task-related variables when the animal transitioned from the A:B to X:X task. Interestingly, neurons encoding chosen juice in the A:B task tended to become untuned in the X:X task. Conversely, neurons encoding Chosen Quantity and Chosen Side in the X:X task tended to become untuned in the A:B task.

The findings thus far suggest the involvement of two dissociable modules in the OFC for representing MC and SC, respectively. One module comprises neurons encoding Offer Value, Chosen Value, and Chosen Juice, while the other module comprises neurons encoding Offer Quantity, Chosen Quantity, and Chosen Side.

To further examine the degree of overlap between these two modules, we conducted the following analysis. In the A:B task, we combined variables Offer value, Chosen value, and Chosen juice into a single variable, MC, representing a multi-attribute choice. Similarly, in the X:X task, we combined variables Offer quantity, Chosen quantity, and Chosen side into a single variable, SC, representing a single-attribute choice. This resulted in two reduced contingency tables: one for the control condition (**Fig. 7a**) and the other for the experimental condition (**Fig. 7b**). As shown in **Fig. 7a**, the cell count for {Symbol #, Symbol #} as well as {MC, MC} was significantly above chance (Symbol #: odds ratio = 12.4, p < 10^-6^, Fisher’s exact test; MC: odds ratio = 1.73, p < 0.05, Fisher’s exact test), indicating consistent representation of Symbol # and MC in the control condition across trial blocks. Intriguingly, the contingency table for the experimental condition (**Fig. 7b**) revealed that when the animal switched from the A:B to X:X task, the likelihood of an MC-representing neuron transitioning to become an SC-representing neuron was significantly below chance (odds ratio = 0.40, p < 10^-4^). Conversely, neurons representing MC in the A:B task tended to become untuned in the X:X task (odds ratio = 2.47, p < 10^-4^, Fisher’s exact test), and neurons representing SC in the X:X task tended to become untuned in the A:B task (odds ratio = 2.58, p < 10^-4^, Fisher’s exact test).

**Figure 7.**
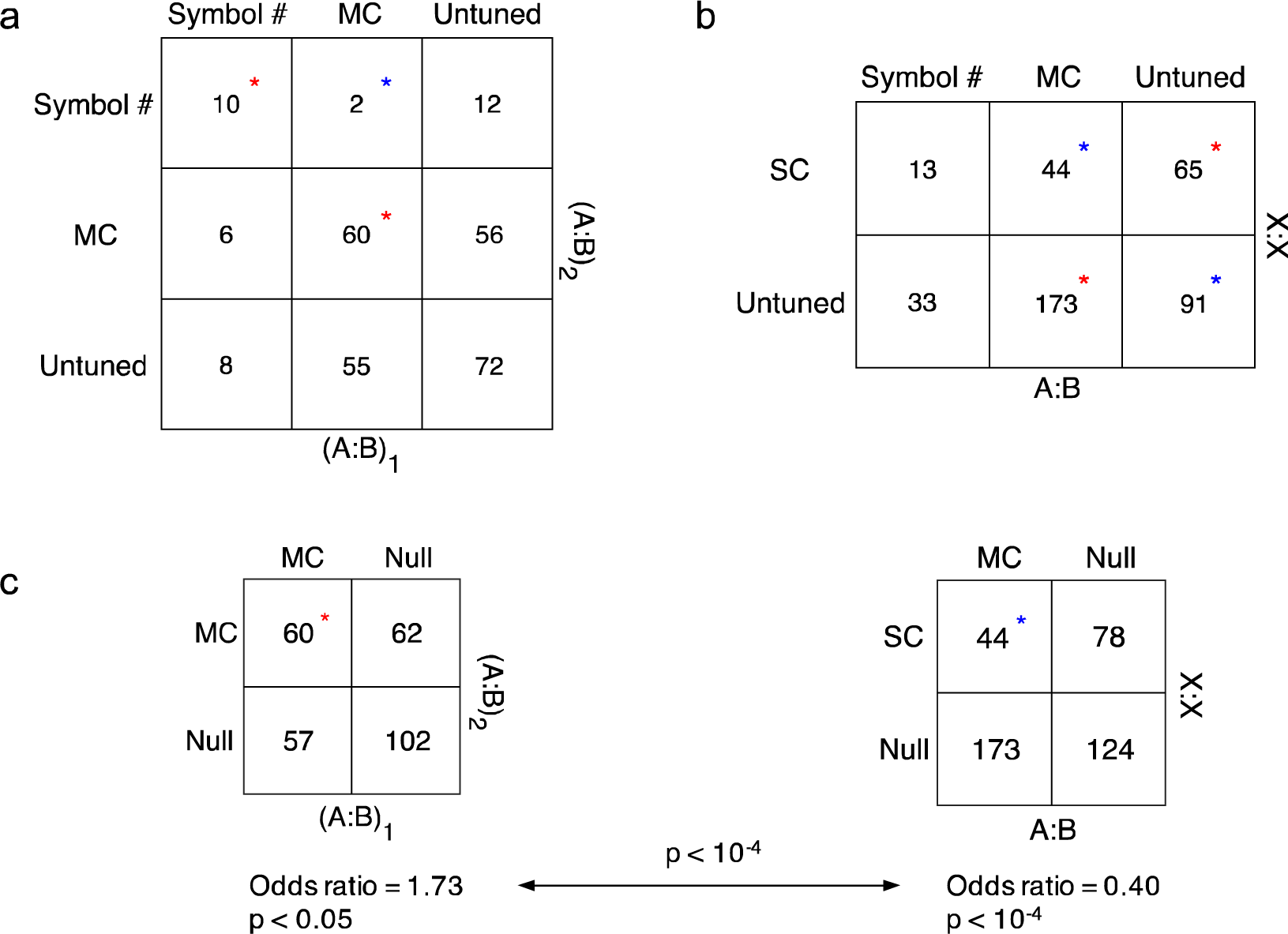
Contrast the remapping between (A:B)_1_→(A:B)_2_ and A:B→X:X conditions. (**a**) Reduced contingency table, (A:B)_1_→(A:B)_2_ condition (n = 281). For each location, we performed Fisher’s exact test. Red asterisks indicate that the cell count was significantly above chance (p < 0.05) while blue asterisk indicates that the cell count was significantly below chance (p < 0.05). MC: neurons encoding variables supporting multi-attribute choice. Task-related cells not explained by any variable were classified as untuned. (**b**) Reduced contingency table, A:B→X:X condition (n = 419). SC: neurons encoding variables supporting single attribute choice. Red asterisks indicate that the cell count was significantly above chance while blue asterisk indicates that the cell count was significantly below chance (p < 0.05). (**c**) Odds ratio for {MC, SC} in A:B→X:X condition is significantly lower than that for {MC, MC} in (A:B)_1_→(A:B)_2_ condition (p < 10^-4^, Z test).

To further investigate how MC and SC are represented during task switching, we compared the contingency table for the control condition (**Fig. 7a**) with that for the experimental condition (**Fig. 7b**). The odds ratio measures the strength of association between two specific variables across the two trial blocks. We observed that the strength of association for {MC, SC} in the A:B → X:X condition is significantly weaker than that for {MC, MC} in the (A:B)_1_ → (A:B)_2_ condition (**Fig. 7c**, p < 10^-4^, Z test).

One caveat in the above analysis is that in the A:B task, the variables Quantity_contra_ and Quantity_ipsi_ are correlated with the variables in the juice-based reference frame. Including Quantity_contra_ and Quantity_ipsi_ in the variable selection analysis may reduce the number of cells that encode variables in the juice-based reference frame, potentially lowering the overlap of cells between MC and SC. To address this, we performed an additional analysis with a conservative approach by excluding Quantity_contra_ and Quantity_ipsi_ in the variable selection analysis for the A:B task.

We found that among the 46 cells that encoded Quantity_contra_ or Quantity_ipsi_, 22 were identified as encoding one of the four variables in the juice-based reference frame, while the remaining 24 did not encode any of the variables and were thus defined as untuned. We then reconstructed the contingency and the odds ratio table (**Fig. S2a** and **S2b**). The general pattern of the table remained the same. Essentially, there was no consistent mapping in the encoding of task-related variables when the animal transitioned from the A:B to X:X task. Meanwhile, neurons encoding chosen juice in the A:B task tended to become untuned in the X:X task. Conversely, neurons encoding Quantity_contra_, Chosen Quantity, and Chosen Side in the X:X task tended to become untuned in the A:B task.

The corresponding reduced contingency table revealed the same trend (**Fig. S2c**). In the experimental condition, when the animal switched from the A:B to X:X task, the likelihood of an MC-representing neuron transitioning to become an SC-representing neuron was even more significantly below chance (odds ratio = 0.34, p < 10^-5^). The strength of association for {MC, SC} in the A:B → X:X condition remained significantly weaker than that for {MC, MC} in the (A:B)_1_ → (A:B)_2_ condition (**Fig. S2c** and **S2d**, p < 10^-4^, Z test).

These findings collectively suggest that dissociable neuronal modules in the OFC represent MC and SC.

### Organization of functional neuronal populations in MC

The aforementioned discoveries suggest that MC is facilitated by a neuronal module dissociable from that supporting SC. Drawing parallels from the neuronal organizations supporting perceptual decision-making, where sensory information is processed by structured neuronal ensembles with organized anatomical connections (Albright et al., 1984; Gottlieb, 2007; Nienborg et al., 2012; Purushothaman and Bradley, 2005), we ask whether similar evidence exists for the structured organization of value-coding neurons in the OFC. While anatomical clustering of functional neuron types in the OFC has not been documented, our study offers a unique opportunity to explore this aspect. This is possible because a majority of the neurons were recorded using 16-channel linear arrays with fixed spacing (150 μm) between adjacent recording channels. Therefore, we utilize the electrode spacing as a surrogate measure for the anatomical distance between pairs of simultaneously recorded neurons.

The distance analysis involved 419 neurons recorded during A:B and X:X tasks, all of which were task-related in at least one of these tasks. Specifically, in the A:B task, 284 neurons were best fit for one of the selected variables: Symbol #_spatial_ (Symbol #_contra_ and Symbol #_ipsi_), Offer value A, Offer value B, Chosen value, or Chosen juice. Pairwise distance analysis was conducted on neuron pairs encoding the same variable: {Symbol #_contra_}_pair_ (14 pairs), {Symbol #_ipsi_}_pair_ (3 pairs), {Offer Value A}_pair_ (19 pairs), {Offer Value B}_pair_ (22 pairs), {Chosen Value}_pair_ (25 pairs), and {Chosen Juice}_pair_ (72 pairs). To establish a null population for the A:B task, an equal number of neuron pairs that were not task-related in either the A:B or X:X tasks were randomly sampled from the trial blocks where at least one encoding pair was recorded. For instance, if in a trial block, there were 2 pairs for {Offer Value B}_pair_ and 3 pairs for {Chosen Juice}_pair_, this block would contribute 5 randomly sampled null pairs to the {Null_A:B_} group, resulting in a total of 155 pairs. In the X:X task, 125 neurons were best fit for one of the selected variables: Offer Quantity_spatial_ (Offer Quantity_contra_ and Offer Quantity_ipsi_), Chosen Quantity, or Chosen side. Neuron pairs encoding the same variable were {Offer Quantity_contra_}_pair_ (11 pairs) and {Chosen Quantity}_pair_ (51 pairs). No neuron pairs were identified for {Offer Quantity_ipsi_}_pair_ or {Chosen Side}_pair_. The {Null_X:X_} group was formed using the same procedure as the {Null_A:B_} group.

In our analysis, we focused on neuron pairs contributing to the putative MC and SC modules. We combined pairs encoding Offer Value A, Offer Value B, and Chosen Value into value coding pairs, labeled as {Value}_pair_. Similarly, pairs encoding Offer Quantity_contra_, Quantity_ipsi_, and Chosen Quantity were grouped as quantity coding pairs, termed {Quantity}_pair_. For each type of coding, we categorized neuron pairs based on whether the encoding slopes of the two cells had the same or opposite signs. For MC, the distance analysis included neuron pairs categorized as {Value}_same_, {Value}_opp_, {Chosen Juice}_same_, {Chosen Juice}_opp_, and {Null_A:B_}. Meanwhile, for SC, the analysis encompassed pairs classified as {Quantity}_same_, {Quantity}_opp_, and {Null_X:X_}.

We compared the distances of neuron pairs encoding the same variable and with the same or opposite encoding signs (**Fig. 8**). In the A:B task, the distances of {Value}_same_ pairs (300 ± 51 μm, median ± ste) were significantly shorter than those for the {Value}_opp_ pairs (900 ± 88 μm) (p < 0.0001, Wilcoxon Rank-Sum Test). Furthermore, the distances of {Value}_same_ pairs were also shorter than those of {Chosen Juice}_same_ pairs (975 ± 101 μm, p < 0.001), {Chosen Juice}_opp_ pairs (600 ± 81 μm, p < 0.01), and {Null_A:B_} pairs (450 ± 40 μm, p < 0.01) (Wilcoxon Rank-Sum Test). In the X:X task, the distances of {Quantity}_same_ pairs (600 ± 101 μm) or {Quantity}_opp_ pairs (1050 ± 81 μm) were not significantly different from each other or from the distances of {Null_X:X_} pairs (750 ± 63 μm) (p > 0.1, Wilcoxon Rank-Sum Test). Across the A:B and X:X tasks, the distances of {Value}_same_ pairs were shorter than those of {Quantity}_same_ or {Quantity}_opp_ pairs (p < 0.01, Wilcoxon Rank-Sum Test).

**Figure 8.**
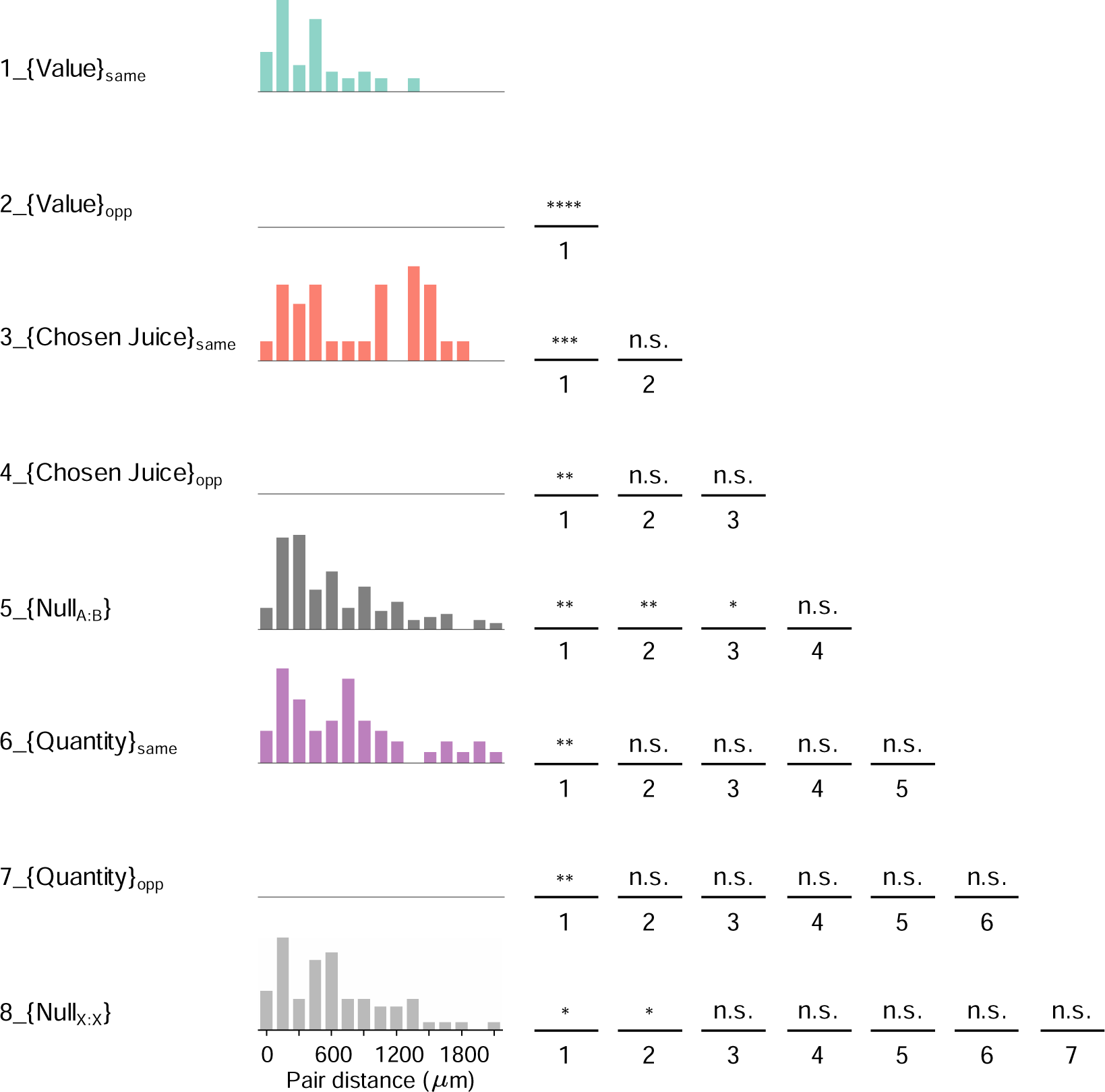
Pair distance for task variable encoding neurons. Distribution of cell pair distance for simultaneously recorded cell pairs encoding the *same* variable. Cells pairs with the same or opposite sign of encoding slopes were grouped separately. Cell pairs encoding Offer Value and Chosen Value in the A:B task were pooled together as {Value}_same_ and {Value}_opp_. Cell pairs encoding Offer Quantity and Chosen Quantity in the X:X task were pooled together as the {Quantity}_same_ and {Quantity}_opp_. {Null_A:B_} consists of randomly sampled cell pairs that were not modulated in either the A:B or X:X task. Elements in the matrix on the right indicate whether the distance between two groups is significantly different from each other. The index of each element corresponds to those indexed for each variable on the ordinate. For example, the element in the top row (****) indicates that the distance of {Value}_same_ is significantly shorter than that for {Value}_opp_. *p < 0.05, **p < 0.01; ***p < 0.001, ****p < 0.0001; n.s., not significant (Wilcoxon Rank-Sum Test).

The results suggest that OFC neurons engaged in value representation in MC tend to be anatomically clustered. However, a potential caveat regarding the outcome in **Fig. 8** is the possibility of false positive results introduced through spike sorting of neurons recorded from the same channel (pair distance = 0). In an extreme hypothetical case, during spike sorting, the spike waveforms of a neuron might be split into two groups, leading to the classification of two different cells. If these “two cells” encode the same value-related variable, there is a higher probability that the encoding slopes have the same rather than the opposite sign, which could erroneously boost the number of zero-distance pairs for the {Value}_same_ group. To address this concern, we adopted a conservative approach by removing all neuron pairs recorded from the same channel (distance = 0). Subsequently, we repeated the analysis performed in **Fig. 8**, and the conclusions remained unchanged (**Fig. S3**).

In summary, in the A:B task, neurons encoding the same value-related variables with the same sign of encoding slopes tend to form anatomical clusters. These findings further support the postulation that the OFC harbors a neuronal module for multi-attribute choice.

## Discussion

The findings of our study provide initial evidence for dissociable neuronal modules within the OFC representing MC and SC tasks. In the MC task, OFC neurons predominantly encoded variables related to offer value, chosen value, and chosen juice, reflecting the abstract representation of subjective value and choice. This aligns with previous studies demonstrating the role of the OFC in encoding multi-attribute choices (Cai and Padoa-Schioppa, 2014; Padoa-Schioppa, 2011; Padoa-Schioppa and Assad, 2006; Pastor-Bernier et al., 2021; Xie and Padoa-Schioppa, 2016). In contrast, in the SC task, where the animal makes choices based solely on the comparison of a single attribute (juice quantity), a separate set of neurons tuned in to encode offer quantity emerged in the OFC. This suggests a shift in neuronal encoding strategies to accommodate the different cognitive demands of the task. Overall, our results highlight the flexibility and adaptability of the OFC in representing different choice tasks. The observed dissociation between MC and SC tasks suggests that the OFC hosts dissociable neuronal modules tailored to specific cognitive processes, such as integrating multiple attributes in MC or focusing on single attributes in SC. These findings contribute to our understanding of the neural mechanisms underlying complex task demands in decision-making and provide insights into how the brain organizes information to support adaptive behavior.

The findings from previous studies by Xie and Padoa-Schioppa (2016) and Shi et al. (2022) provide compelling evidence for the adaptability of the neuronal circuitry underlying economic choice in the OFC. In Xie and Padoa-Schioppa’s study, the persistence of neuronal encoding across different pairs of juices suggests that the same neural circuitry can flexibly accommodate varying sensory inputs without fundamentally altering its encoding identity. Similarly, Shi et al. (2022) demonstrated that the same neural circuit can support economic choices with different temporal task structures, further highlighting the adaptive nature of OFC neuronal encoding. Building upon these previous findings, our current study extends the understanding of OFC neuronal organization by revealing dissociated modules dedicated to different choice tasks. While the OFC neurons are capable of adapting their encoding identity across different task contexts, as demonstrated by Xie and Padoa-Schioppa (2016), our results suggest that there is likely a dedicated mapping between choice tasks and the underlying neuronal modules.

The clustering of value-coding neurons in the OFC, as demonstrated in our study, offers intriguing insights into the organization and functional significance of this brain region in economic decision-making. There are several possible explanations for the observed clustering phenomenon. Firstly, the anatomical clustering of value-coding neurons in the OFC may have developmental significance. The OFC receives inputs from various sensory and limbic regions, integrating information related to reward, emotion, and sensory perception (Ongur and Price, 2000). The clustering of value-coding neurons could emerge as a result of the developmental wiring of these converging inputs. Given that connecting distant neurons requires energy-intensive long-distance connections, clustering may serve to minimize the biological cost associated with wiring, facilitating efficient information processing (Chklovskii and Koulakov, 2004). Alternatively, the observed clustering of value-coding neurons could arise from extensive experience in performing the juice choice task. Similar to the formation of category-selective regions in the macaque temporal lobe during symbol training (Srihasam et al., 2012), repeated exposure to specific decision-making tasks may lead to the organization of neurons into functional clusters optimized for processing relevant information. In this scenario, the clustering reflects the plasticity of the OFC in adapting to task demands and optimizing neural processing efficiency. Whether driven by developmental wiring, or task-related experience, the clustering phenomenon underscores the importance of value processing in shaping the functional architecture of the OFC.

The existence of dissociable neuronal modules for MC and SC raises intriguing questions about the functional organization of the. Several postulations may shed light on why these two modules are dissociable and how they may contribute to decision-making processes. Firstly, the time required for restructuring the same neuronal module after each task switch may exceed the duration of a single experimental session in our study. Neuronal circuits in the brain are highly dynamic and can undergo rapid reorganization in response to changing task demands. However, significant restructuring of neural circuits may require time and repeated exposure to new task conditions. Therefore, maintaining two dissociable modules configured for each task allows for efficient task performance without the need for extensive neural reorganization between trials. Secondly, restructuring neuronal modules may incur significant metabolic and computational costs. Maintaining separate modules for MC and SC may be advantageous from an energy efficiency standpoint, as it minimizes the need for frequent neural reconfiguration and conserves metabolic resources.

The third and perhaps most plausible possibility is that the dissociable modules observed in the OFC represent an established circuit for economic choice. Analogous to face patches in the inferotemporal cortex for face processing (Tsao et al., 2006), these functional groups of neurons in the OFC may constitute specialized circuits that have evolved to support specific aspects of decision-making. Such organized circuits may not only provide an abstract cognitive map for the flexible organization of goods information but also serve as an anatomical substrate underlying the transitivity of preferences, a fundamental concept in rational choice behavior (Cai, 2021; Padoa-Schioppa and Assad, 2008; Rustichini et al., 2017). By maintaining dissociable modules for MC and SC, the brain may optimize decision-making processes by efficiently leveraging specialized circuits tailored to different task demands.

## Methods

### Experimental design and neuronal recordings

Two adult male rhesus monkeys (*Macaca mulatta*; monkey S, 11.4 kg; monkey N, 12.2 kg) participated in the study. All experimental procedures conformed to the NIH *Guide for the Care and Use of Laboratory Animals* and were approved by the Institutional Animal Care and Use Committee at NYU Shanghai. Before training, a head restraining device and an oval recording chamber (50 × 30 mm) were implanted on the skull under general anesthesia. The chamber allowed access to bilateral OFC with penetrations on a nearly coronal plane. During the experiments, monkeys sat in an electrically insulated recording room with their heads restrained. Visual stimuli were presented on a computer monitor placed 57 cm in front of the monkey. Eye positions were monitored with an infrared video camera (Eyelink, SR research). The behavioral task was controlled through custom-written software (http://www.monkeylogic.net) based on Matlab (MathWorks).

Structural MRI scans were obtained before and after the surgery. In the post-surgery MRI, a custom-made plastic insert was placed in the recording chamber with a tight fit. Cylindrical channels in the inset parallel to the path of the electrodes were filled with vitamin E oil for imaging. Coronal images were reconstructed from the MRIs to provide a reference for recording locations. Neuronal data were collected from one hemisphere of monkey S and both hemispheres of monkey N (**Fig. 2**). Most of the recordings were performed using one or two 16-channel linear arrays (U- or V-probes; Plexon). U/V-probes had a diameter of 185 μm and contacts were 150 μm apart. The electrodes, placed ≥1 mm apart from each other, were advanced using a motorized micro-drive system (THOMAS RECORDING). A subset of data was collected using individual tungsten electrodes (125 μm diameter, Frederick Haer). Tungsten electrodes were advanced with the same micro-drive system used for linear arrays. After the electrodes reached the target sites in the OFC, we waited about 20 minutes for the tissue to settle before starting each recording session to promote signal stability. Electrical signals were recorded using a 32-channel system (Plexon Inc). Action potentials were detected online and saved to disk for spike sorting using standard software (Offline Sorter, Plexon Inc). Only neurons that remained stable and well-isolated during two consecutive trial blocks were included in the analysis.

**Fig. 1a** illustrates the experimental design. In each trial, the animal chose between two juices offered in variable amounts. The trial began with the monkey gazing a fixation point in the center of the monitor (center fixation window: 3°). After 0.5 s, two sets of squares representing the two offers appeared on the two sides of the fixation point. For each offer, the color indicated the juice type and the number of squares indicated juice quantity, with each square representing a juice quantum 100 μl). The animal maintained center fixation for a randomly variable delay (0.6–1.2 s, uniform distribution) followed by a go signal, which was indicated by the extinction of the center fixation point and the appearance of two saccade targets. The monkey had 1.5 s to indicate its choice with a saccade and was required to maintain target fixation (peripheral fixation window: 4°) for 0.3 s before juice delivery.

In our study, A:B indicates choice between two juices with A as the one with the preferred taste; B:B or A:A indicates choice between the same juice. The same two juices were used in each session. Since each session included two A:B blocks, they are named in their temporal sequence in the experimental session. For monkey S, the block sequence is B:B, (A:B)_1_, (A:B)_2_, A:A; For monkey N, the block sequence is B:B, (A:B)_1_, (A:B)_2_, A:A or (A:B)_1_, (A:B)_2_, A:A, (A:B)_3_. Based on our design, (A:B)_1_→ (A:B)_2_ serves as the control condition while (A:B)_1_→ B:B and (A:B)_2_→ A:A serve as the contrast for neuronal representation in MC and SC and are collectively termed A:B→X:X.

The offer combinations for the A:B blocks arranged in descending order of the (A/B) quantity ratio is: [1A:0B 3A:1B 2A:1B 2A:2B 1A:2B 1A:3B 1A:4B 1A:6B 0A:2B]. To perform the matching analysis as that for the A:B block, in B:B block, in which the same juice was used, one of the two offers was coded as lowercase “a”. Thus, the offer combination for B:B blocks was coded as [2a:0B 6a:1B 4a:1B 3a:1B 2a:1B 2a:2B 1a:2B 1a:3B 1a:4B 1a:6B 0a:2B]. Similarly, the offer combination in the A:A block was coded as: [2A:0b 3A:1b 2A:1b 1A:1b 1A:2b 1A:3b 0A:2b]. The offer combinations for B:B and A:A blocks are designed such that the value range for offer A and offer B remains the same as that in the A:B block. For the A:B task, a “trial type” was defined by two offers and a choice (e.g., [1A+:3B-, A]). In this notation, “+” indicates offer located on the left hemifield while “-” indicates the offer located on the right hemifield. For the B:B or A:A task, because therefore is only one type of juice and the animals almost always choose the larger quantity offer, “trial type” was defined by two offers and the location of the larger quantity offer (e.g., [1B+:3B-]).

Blinding was not used in the behavioral design. In each block, offer types were pseudo-randomly interleaved and the left/right configurations of the offers were counterbalanced. In all sessions, the two blocks were separated by a ∼5 min break. The number of cells collected for each condition and the number of trials run for each cell were not predetermined using a statistical method but were comparable to those of previous studies.

### Behavioral analysis and neuronal classification

Behavioral data were analyzed separately for each trial block. The ‘choice pattern’ was defined as the percent of trials in which the animal chose the non-preferred juice as a function of the log quantity ratio of the two offers. In the analysis, we fitted the choice pattern with a normal cumulative distribution function. The flex of the sigmoid, corresponding to the indifference point (ρ), provided a measure for the relative value between the two offers.

The analysis performed for each cell in each trial block followed the procedures used in previous studies. Neuronal activity was examined in two time windows: pre-offer (0.5 s before offer onset), and post-offer (0.1∼0.6 s after the offer onset) window. Firing rates were obtained by averaging spike counts over each time window and across trials for each trial type. A ‘neuronal response’ was defined as the firing rate of one cell, in one block, in one time window, as a function of trial type. The two responses recorded for the same cell, in the same time window, in the adjacent two blocks is defined as a ‘response pair’. For each neuron and each time window, we performed a one-way ANOVA (factor: trial type) separately in each block. We imposed a p < 0.01 threshold and admitted to subsequent analyses only response pairs that met this criterion in at least one block. Neurons that passed this ANOVA criterion in at least one time window were identified as ‘task-related’.

Data from each trial block were considered separately. The entire data set for the A:B task included 2,029 cells. Previous work showed that the vast majority of neuronal responses encoded one of three variables: Offer Value, Chosen Value, and Chosen Juice. For each variable, the encoding was linear and either positive (increasing firing rate for increasing values) or negative (increasing firing rate for decreasing values). Thus, as a preliminary step, we repeated the variable selection analyses conducted previously on the current data set in the A:B task. We computed 11 variables (Table S1) and applied the stepwise and best-subset procedures for variable selection. A variable was said to ‘explain’ the response if the regression slope differed significantly from zero (p < 0.05). The regression also provided an *R*^2^, which was set equal to zero if the variable did not explain the response. The variable with the highest *R*^2^ is assigned to the cell as the encoded variable. Both procedures selected variables Symbol #_spatial_, Offer value_juice_, Chosen Value, and Chosen Juice. The same analysis was applied to the X:X task (1,964 cells) with another set of variables potentially encoded (Table S1) and the selected variables are Offer Quantity_spatial_, Chosen Quantity, and Chosen Side.

### Comparing neuronal classifications across tasks

To compare the classifications across trial blocks at the population level, we constructed contingency tables separately for data collected in (A:B)_1_→ (A:B)_2_ condition and A:B→X:X condition. The rows and columns of the tables represent the classifications of neurons in the two blocks respectively, while the numbers represent cell counts. Our initial analysis was to assess, for each location in the contingency table, whether the cell count deviated significantly from chance level, assuming independent classifications in the two trial blocks. To this end, we used the statistical test based on odds ratio.

We indicate with *X_i,j_* the number of cells encoding variable *i* in the first block and variable *j* in the second block. For each location (*i*, *j*) in the table, we computed a 2 × 2 matrix with elements

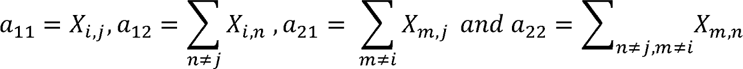

The odds ratio for location (*i*, *j*) was defined as OR*_i,j_* = (*a*_11_/*a*_12_)/(*a*_21_/*a*_22_). If the likelihood of a neuron encoding variable *i* in the first block is independent of the likelihood of it encoding variable *j* in the second block, the expected value of OR*_i_*_,*j*_ equals 1. In other words, the chance level for odds ratio is 1. Conversely, OR*_i,j_* > 1 (OR*_i,j_*< 1) indicated that the cell count in location (*i*, *j*) was above (below) chance level. To assess whether departures from chance level were statistically significant, we used Fisher’s exact test (two tails).

### Comparing the remapping across the control and experimental condition

In a series of analyses, we directly compared the classification results obtained for the control ((A:B)_1_→ (A:B)_2_) and experimental (A:B→X:X) conditions. For each condition, we constructed a reduced contingency table. We then constructed 2 × 2 contingency tables for the elements of interest. Finally, we compared the odds ratios associated with each corresponding pair of elements with a Z-test.

### Verification of recording stability across trial blocks

The validity of the comparison of cross-block representations for (A:B)_1_→ (A:B)_2_ and A:B→X:X conditions is contingent upon the assumption that recording was stable across two trial blocks and there is no significant difference between the stability of recording across the above two conditions.

To test this assumption, we computed the Pearson correlation for both the distribution of inter-spike intervals (CC_isi_same_) and the spike waveforms (CC_waveform_same_) of the same neuron across the two adjacent blocks. We also constructed a dataset of control distribution as CC_isi_random_ and CC_waveform_random_ by pairing a neuron in one block with a randomly selected different neuron in the next block. Finally, we computed the difference of the Pearson correlation between the experiment and control condition as the recording stability index (RSI) based on inter-spike intervals and the spike waveforms as follows.

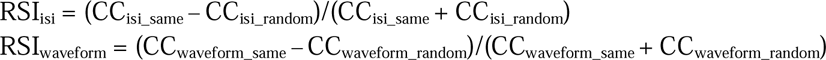

Lastly, we compared the RSI_isi_ and RSI_waveform_ between (A:B)_1_→ (A:B)_2_ and A:B→X:X conditions using the Wilcoxon Rank-Sum Test.

### Analysis of neuron pair distance

In our study, most of the neurons were recorded with 16-channel linear arrays with known spacing (150 μm) between adjacent recording channels. Therefore, we utilize the electrode spacing as a surrogate measure for the anatomical distance between pairs of simultaneously recorded neurons. If a pair of neurons were recorded from the same channel, their distance was considered zero.

## Acknowledgments

We thank Yuanyuan Liu for help with animal training and Camillo Padoa-Schioppa and Weikang Shi for helpful comments on the manuscript. This research was supported by the National Science and Technology Innovation 2030 Major Program (No. 2021ZD0203700/2021ZD0203702 to X.C.) and Shanghai Municipal Science and Technology Major Project (2018SHZDZX05 to X.C.).

## Data availability

The datasets generated during and/or analyzed during the current study are available from the corresponding author upon reasonable request.

## Code availability

All analyses were performed using custom code written in Matlab (MathWorks) and available upon request.

---

**Table S1.** Defined variables. For the A:B task, in any given trial, q_A_ and q_B_ were, respectively, the quantities of juice A and juice B offered to the animal, r was the relative value of the two juices obtained from a sigmoid fit.

| **A:B** | **Collapsed variable** | **Variable** | **Definition** | **Reference frame** |
| --- | --- | --- | --- | --- |
| 1 | Offer Value (juice) | Offer Value A | $p q_A$ | Juice |
| 2 | | Offer Value B | $q_B$ | Juice |
| 3 |  | Chosen Juice | 1 if juice B is chosen, 0 if juice A is chosen | Juice |
| 4 |  | Chosen Value | offer value A if juice A chosen, offer value B if juice B chosen |  |
| 5 |  | Other Value | offer value B if juice A chosen, offer value A if juice B chosen |  |
| 6 | Offer Value (spatial) | Offer Value<sub>contra</sub> | offer value A/offer value B if offer A/offer B is on the contralateral hemifield | Action |
| 7 |  | Offer Value<sub>ipsi</sub> | offer value A/offer value B if offer A/offer B is on the ipsilateral hemifield | Action |
| 8 |  | Chosen Side | 1 if contralateral offer is chosen, 0 if ipsilateral offer is chosen | Action |
| 9 | Symbol # (spatial) | Symbol #<sub>contra</sub> | number of squares associated with the offer in the contralateral hemifield | Visual |
| 10 |  | Symbol #<sub>ipsi</sub> | number of squares associated with the offer in the ipsilateral hemifield | Visual |
| 11 |  | Position of A | 1 if offer A is on the contralateral hemifield, 0 if offer A is on the ipsilateral hemifield |  |
| **X:X** |  |  |  |  |
| 1 | Offer Quantity (spatial) | Offer Quantity<sub>contra</sub> | quantity of the juice offered on the contralateral hemifield | Action |
| 2 |  | Offer Quantity<sub>ipsi</sub> | quantity of the juice offered on the ipsilateral hemifield | Action |
| 3 |  | Chosen Side | 1 if contralateral offer is chosen, 0 if ipsilateral offer is chosen | Action |
| 4 |  | Chosen Quantity | quantity of the chosen offer |  |
| 5 |  | Other Quantity | quantity of the unchosen offer |  |

**Figure S1.**
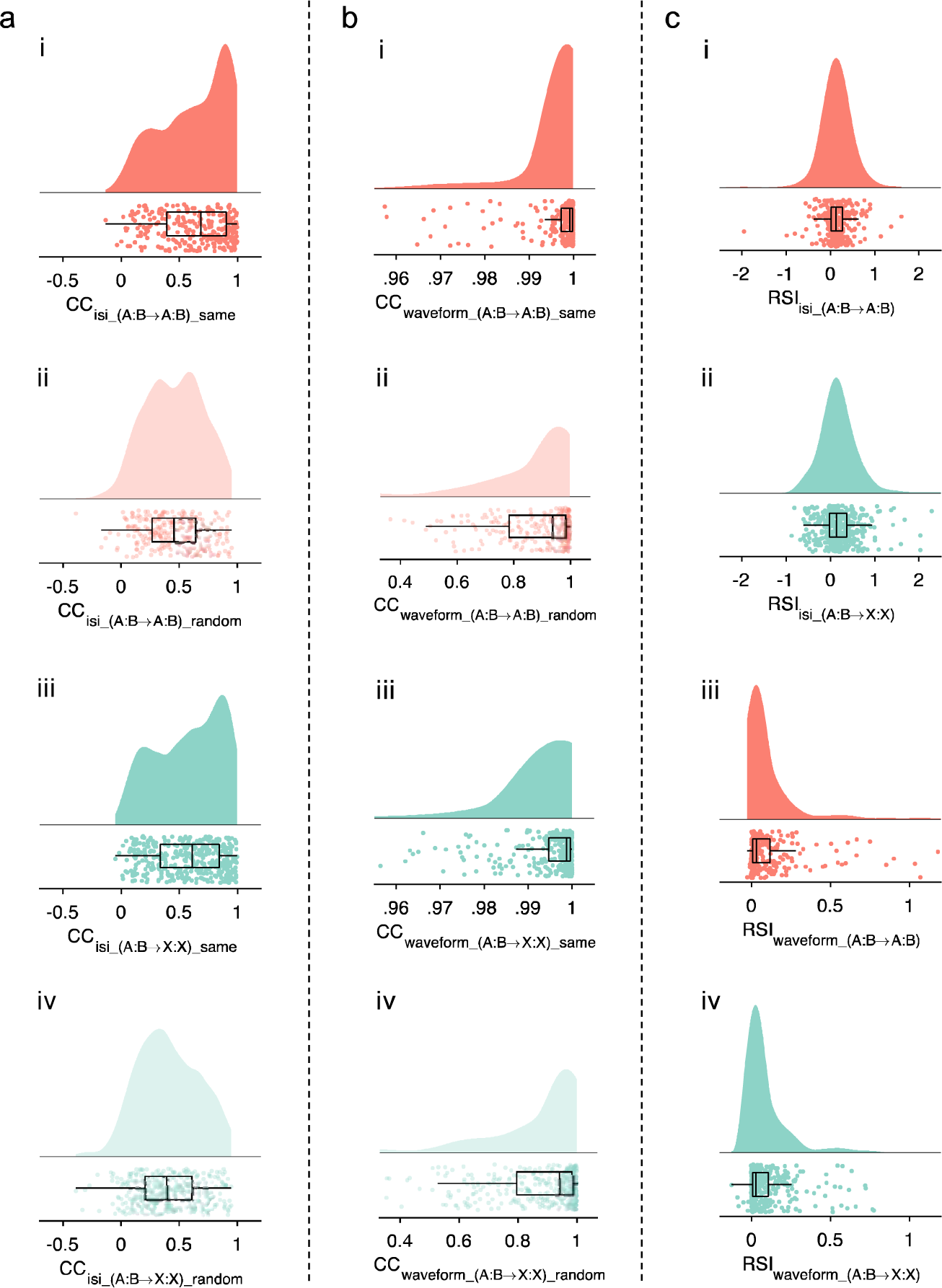
Metrics of recording stability. (**a**) Distribution of Pearson correlation for inter-spike intervals. (**a_i**) Distribution for the same neuron in (A:B)_1_→(A:B)_2_ condition. (**a_ii**) Distribution for the same neuron in A:B→X:X condition. (**a_iii**) Distribution for two randomly paired different neurons in (A:B)_1_→(A:B)_2_ condition. (**a_iv**) Distribution for two randomly paired different neurons in A:B→X:X condition. (**b**) Distribution of Pearson correlation for spike waveforms. (**b_i**) Distribution for the same neuron in (A:B)_1_→(A:B)_2_ condition. (**b_ii**) Distribution for the same neuron in A:B→X:X condition. (**b_iii**) Distribution for two randomly paired different neurons in (A:B)_1_→(A:B)_2_ condition. (**b_iv**) Distribution for two randomly paired different neurons in A:B→X:X condition. (**c**) Distribution of recording stability index. (**c_i**) Distribution of RSI_isi_ for (A:B)_1_→(A:B)_2_ condition. (**c_ii**) Distribution of RSI_waveform_ for A:B→X:X condition. There is no significant difference for RSI_isi_ between (A:B)_1_→(A:B)_2_ and A:B→X:X conditions (p = 0.49, Wilcoxon Rank-Sum Test). (**c_iii**) Distribution of RSI_waveform_ for (A:B)_1_→(A:B)_2_ condition. (**c_iv**) Distribution of RSI_waveform_ for A:B→X:X condition. There is no significant difference for RSI_waveform_ between (A:B)_1_→(A:B)_2_ and A:B→X:X conditions (p = 0.66, Wilcoxon Rank-Sum Test).

**Figure S2.**
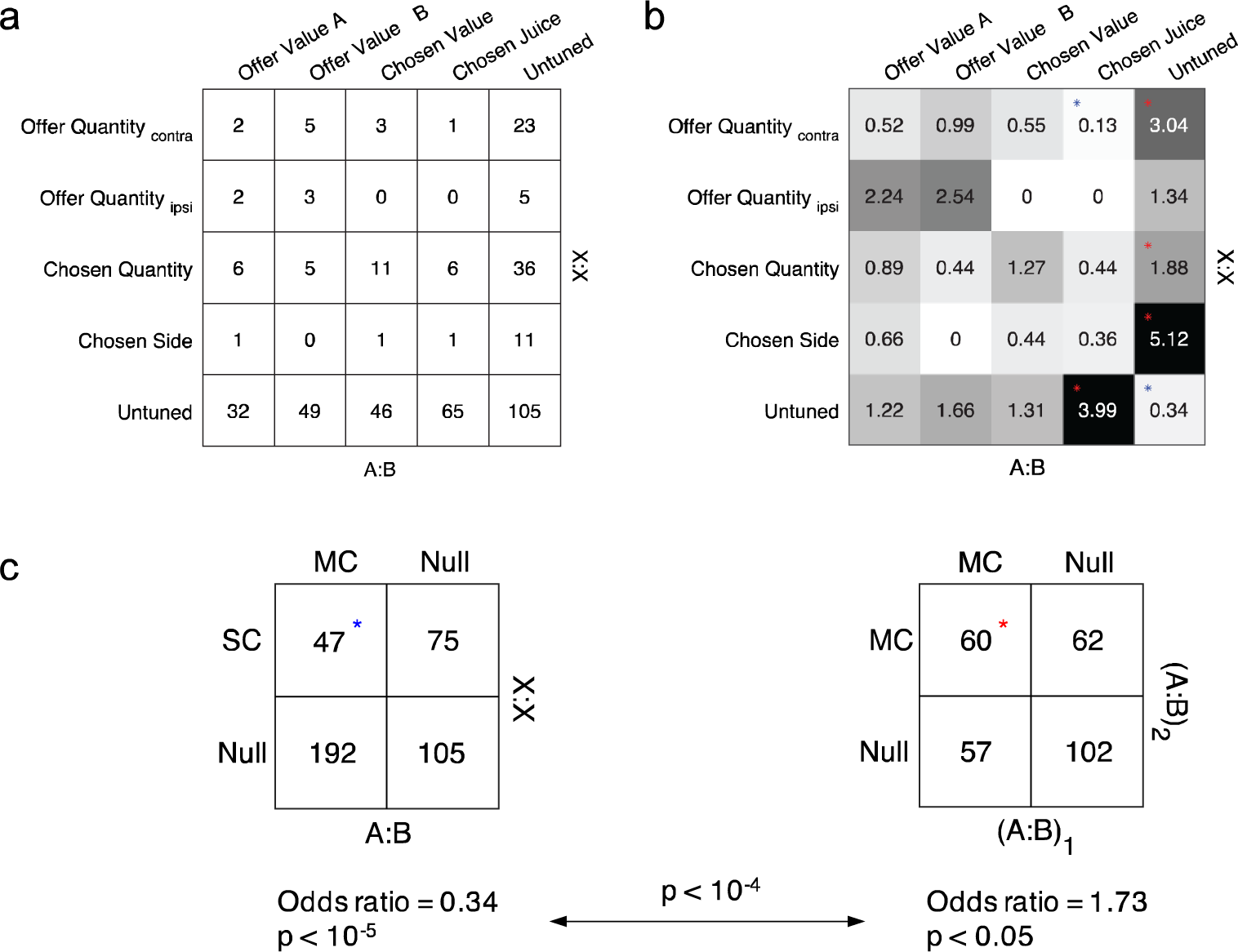
Remapping analysis for A:B→X:X, excluding Quantity_contra_ and Quantity_ipsi_ in the variable selection analysis for the A:B task. (**a**) Contingency table (*n* = 419). Rows and columns represent, respectively, the classification obtained in the A:B and the adjacent X:X blocks. Numbers in the table indicate cell counts. Task-related cells not explained by any variable were classified as untuned. (**b**) Analysis of odds ratios. Numbers in the table and grayscale colors represent the odds ratios obtained for each location in (**a)**. The chance level is 1 and numbers >1 (or <1) indicate that the cell count was above (or below) that expected by chance. For each location, we performed Fisher’s exact test. Red asterisks indicate that the cell count was significantly above chance (p < 0.05) while blue asterisks indicate that the cell count was significantly below chance (p < 0.05). (**c**) Odds ratio for {MC, SC} in A:B→X:X condition is significantly lower than that for {MC, MC} in (A:B)_1_→(A:B)_2_ condition (p < 10^-4^, Z test).

**Figure S3.**
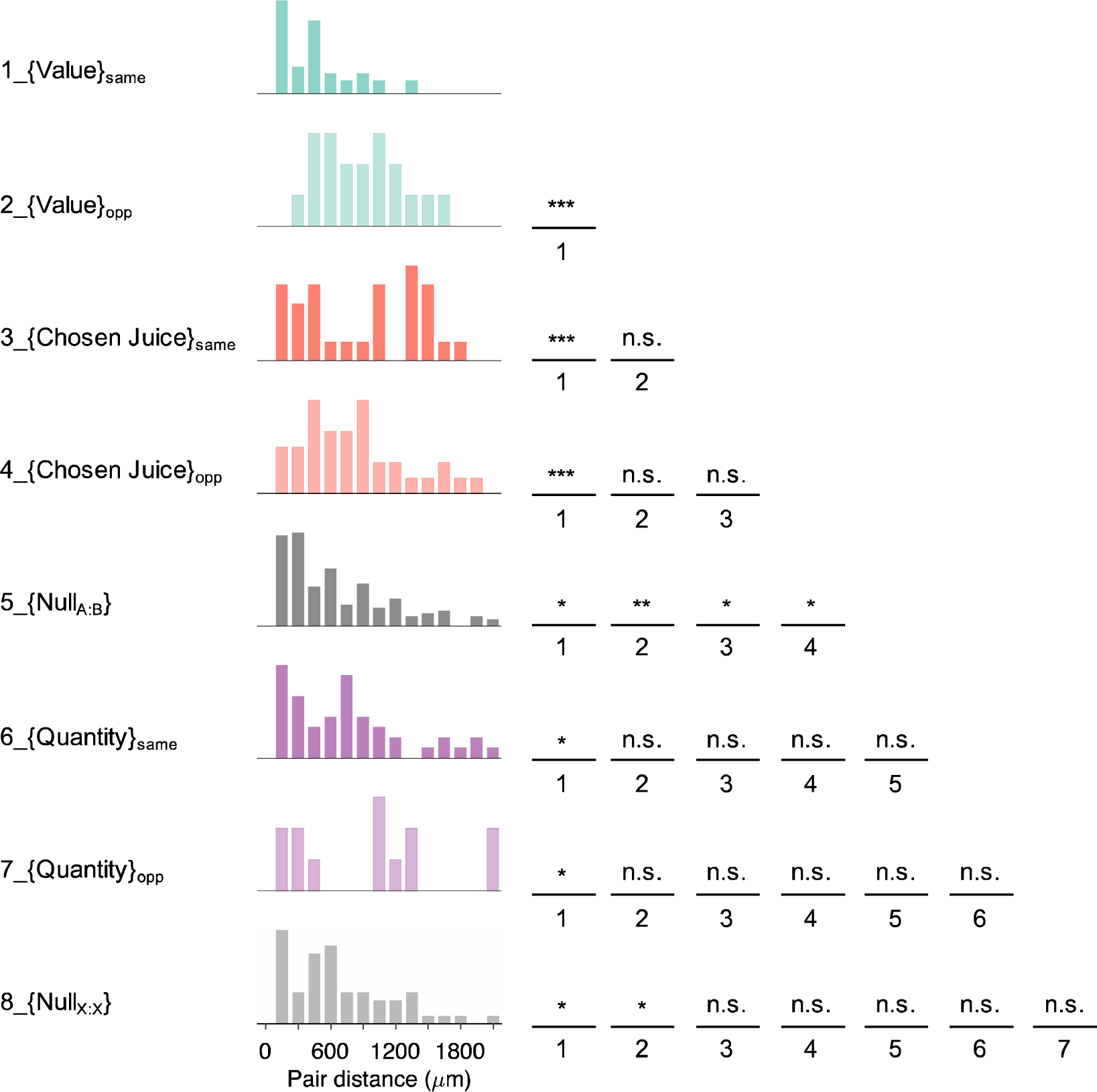
Same as that in Fig. 8 but removing neuron pairs recorded from the same channel (distance = 0). *p < 0.05, **p < 0.01, ***p < 0.001; n.s., not significant.

